# Direct reprogramming of astrocytes to neurons leads to functional recovery after stroke

**DOI:** 10.1101/2020.02.02.929091

**Authors:** Jessica Livingston, Tina Lee, Emerson Daniele, Clara Phillips, Alexandra Krassikova, Tom Enbar, Ines Kortebi, K.W. Annie Bang, Brennan Donville, Omadyor Ibragimov, Nadia Sachewsky, Cindi M Morshead, Maryam Faiz

**Affiliations:** Department of Surgery, University of Toronto; Donnelly Center, University of Toronto; Lunenfeld-Tanenbaum Research Institute, Flow Cytometry Facility, University of Toronto

**Keywords:** reprogramming, direct lineage conversion, NeuroD1, brain repair, stroke, functional recovery

## Abstract

Stroke is the leading cause of adult disability with few treatment options for stroke survivors. Astrocyte reprogramming to neurons enables the targeted *in vivo* generation of new cells at the site of injury and represents a novel approach for brain repair. A number of studies have demonstrated successful conversion of astrocytes to neurons in various models of brain injury and disease; however, the impact of this strategy on tissue and functional outcome following stroke is not well established. Using AAV delivery of the transcription factor NeuroD1, we reprogrammed astrocytes 7 days after endothelin-1 induced cortical stroke, and studied the long-term cellular and functional outcomes. We found that by 63 days post-stroke, 20% of neurons in the perilesional cortex were reprogrammed. Furthermore, reprogrammed neurons had matured into regionally appropriate neuronal subtypes. Importantly, this treatment was associated with improved functional outcome using the foot fault test and gait analysis. Together, our findings indicate that *in vivo* reprogramming is a promising regenerative approach for stroke repair.

## Introduction

Direct lineage reprogramming is emerging as a new approach to replace lost cells for brain repair. Since the seminal discovery that somatic cells can be reprogrammed to induced pluripotent stem cells [1], it has been established that cells can be directly converted from one mature cell type to another [2–6]. Reprogramming strategies have typically used induced expression of transcription factors (TFs) known to act as fate determinants of the target cell during embryonic development [2–4,6] to generate reprogrammed cells both *in vitro* and *in vivo*.

In the central nervous system (CNS) direct lineage reprogramming of astroglial cells to neurons has been demonstrated using different TF combinations that can direct the acquisition of regionally appropriate [7]and functional neuronal subtype identities (e.g.; glutamatergic, GABAergic) [8–11]. Of particular interest in the context of CNS injury is the application of direct lineage reprogramming *in vivo*. Recent work by a number of groups have validated the feasibility of astrocyte-to-neuron conversion in pre-clinical models of Alzheimer’s Disease, Parkinson’s Disease, stab wound injury and most recently, stroke [12–17]. Typically, these studies have used single neural-specific bHLH TFs including Ascl1, Neurog2 and Neurod1, with varying degrees of reprogramming efficiency reported [13,14,16]. However, a considerable gap exists in terms of understanding the behavioural outcomes of astrocyte-to-neuron reprogramming. One study demonstrated that astrocyte conversion to dopaminergic neurons improves some aspects of motor behavior in a mouse model of Parkinson’s Disease [12], and more recently, a report demonstrating improved functional outcomes following astrocyte-to-neuron conversion following ischemia has highlighted the promising potential of cellular reprogramming for functional recovery [17].

Herein, we used adeno-associated viral (AAV) delivery of the TF *NeuroD1* along with cell lineage tracking approaches to characterize astrocyte-to-neuron reprogramming and examine tissue and functional outcomes in a preclinical model of cortical ischemic stroke that replicates the pathophysiological processes of human stroke.

We show that *NeuroD1*-induced conversion of perilesional astrocytes to neurons following stroke leads to the generation of regionally appropriate cortical neurons. Importantly, we demonstrate that astrocyte-to-neuron reprogramming leads to improved motor outcomes, thereby demonstrating that direct cellular reprogramming represents an exciting and promising strategy to treat the stroke injured brain.

## Results

### Delivery of NeuroD1 in the injured cortex leads to astrocyte-to-neuron reprogramming

We designed a reprogramming strategy to permanently label astrocytes after delivery of *NeuroD1*, allowing us to track cell fate during the reprogramming process. Cre conditional YFP reporter mice (YFP^fl^) were transduced with AAV5 that delivered *NeuroD1* linked to Cre under the control of the astrocyte promoter GFAP (AAV5-*NeuroD1*). One week following stroke (post-stroke day 7; PSD 7) AAV5-*NeuroD1* or a Cre control (AAV5-Cre) was delivered to the injured cortex of YFP^fl^ mice (Figure 1a). To determine the efficiency of viral transduction, we isolated GLAST+ astrocytes from the stroke-injured cortex 3 days post-AAV injection (PSD 10) and analyzed the number of YFP+ cells within this population using ImageStream. We found that 50% of the perilesional GLAST+ astrocytes expressed YFP (Figure 1b). Next, we determined the extent of astrocyte-to-neuron reprogramming of the transduced cells. We analyzed the number of reprogrammed neurons (NeuN+YFP+) as a percentage of the total number of YFP+ (transduced) cells at PSD 28. Reprogrammed neurons were found in the perilesional stroke-injured cortex (Figure 1c) and resulted in a 27% reprogramming efficiency 21 days post AAV injection (Stroke_AAV5-*NeuroD1*, 26.73 ± 4.57 %; Stroke_AAV-Cre, 3.37 ± 0.37 %; p=0.036).

**Fig. 1.**
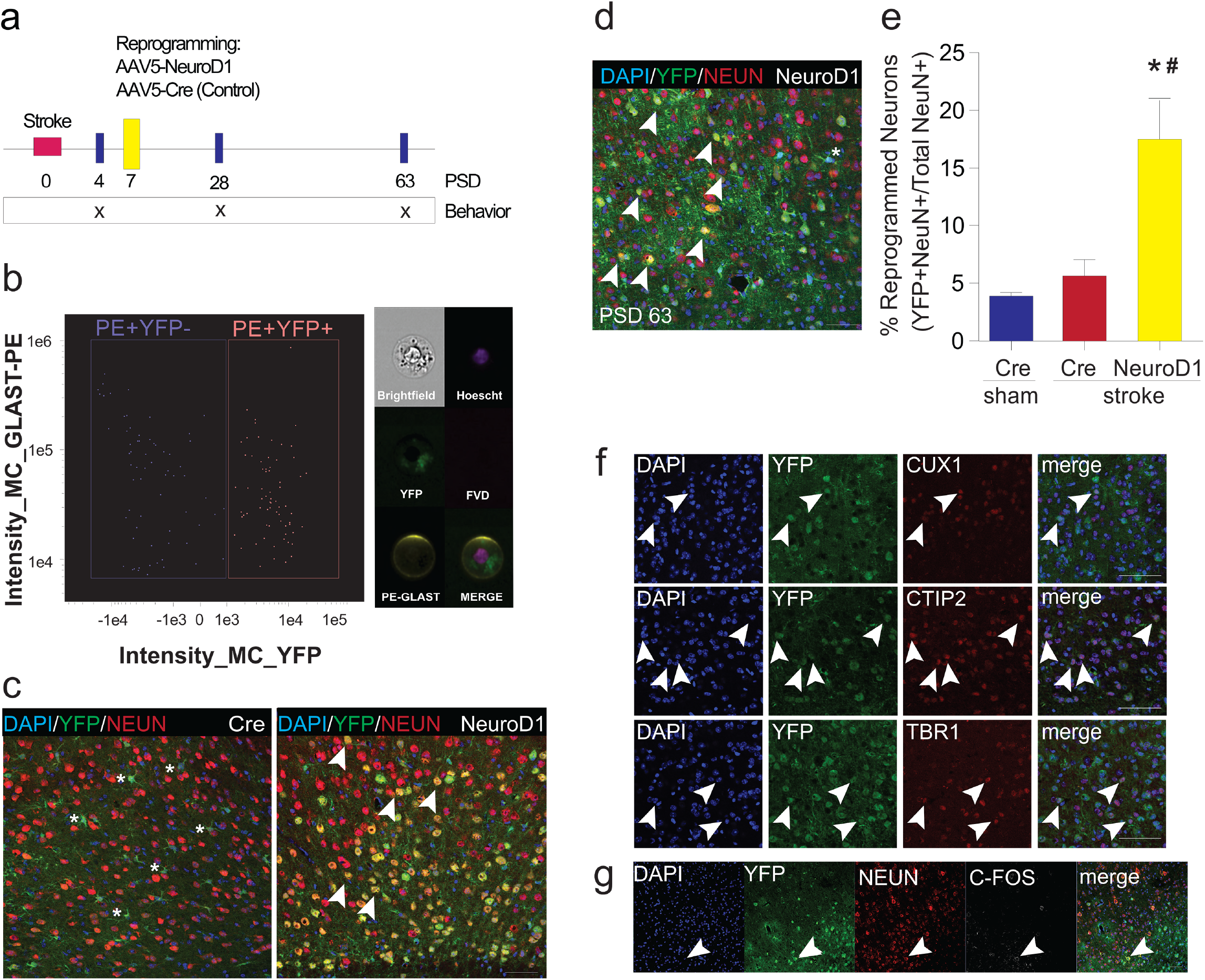
Astrocyte-to-neuron reprogramming after stroke. a) Experimental design. b) ImageStream plot of MACS sorted astrocytes. Pictures depict a single (brightfield), live [fixable viability dye (FVD-)], nucleated (DAPI+), PE+YFP+ astrocyte. c) YFP+NeuN- (asterisks) transduced cells were seen at PSD 28 in brains from Stroke_AAV5-Cre treated mice. Brains from Stroke_AAV5-*NeuroD1* mice contained YFP+/NeuN+ reprogrammed neurons (arrowheads; n=3/group). d) YFP+NeuN+ reprogrammed neurons (arrowheads) at PSD 63 after delivery of AAV5-*NeuroD1*. Asterisk is YFP+/NeuN-astrocyte e) Percent contribution of reprogrammed neurons to the stroke-injured area (n=18 Stroke_AAV5-*NeuroD1;* n=6 Stroke_AAV5-Cre; n=6 Sham_AAV5-Cre; *p<0.05 compared to Sham_AAV5-Cre, #p<0.01 compared to Stroke_AAV5-Cre). f) CUX1+ upper layer and CTIP2+TBR1+ lower layer YFP+ reprogrammed neurons. Arrowheads indicate examples of double-labeled neurons. g) c-FOS expression in NeuN+YFP+reprogrammed neurons (arrowheads). Scale bars = 50 μm. Data are expressed as mean ± SEM.

### Reprogrammed neurons persist long-term

To determine whether reprogrammed neurons persist long-term, we quantified NeuN+YFP+ reprogrammed neurons in the injured cortex at PSD 63 (56 days post-AAV injection). We found approximately 20% of all neurons in the stroke-injured cortex were reprogrammed, which was greater than Stroke_AAV5-Cre and Sham_AAV5-Cre groups (Stroke_AAV5-*NeuroD1*, 18.55 +/- 2.74 %;Stroke_AAV-Cre, 5.66 +/- 1.40 %; Sham_AAV5-Cre 3.91 +/- 0.30 %; p<0.05; Figure 1d-e).

### Reprogrammed neurons differentiate into regionally appropriate cell types

The cortex is a six-layered structure comprised of distinct populations of projection neurons that are located in different cortical layers and exhibit unique morphologies, projection patterns, and gene expression^18^. Using TFs Nurr1 and Ngn2, a recent study showed that astrocyte reprogramming results in regionally appropriate upper and lower layer cortical neurons. To determine whether *NeuroD1* reprogrammed neurons mature into regionally appropriate cortical phenotypes following stroke, we looked for the presence of NeuN+YFP+ neurons that co-expressed markers of upper- and lower-layer cortical neurons. Strikingly, CUX1, a TF present in layer 2/3 neurons, was exclusively expressed in NeuN+YFP+ cells in the upper layers. Similarly, TBR1 and CTIP2, TFs expressed in layer 5/6 neurons, were exclusively expressed in NeuN+YFP+ in the lower layers (Figure 1f). We also examined c-FOS expression, an early immediate gene transcribed during neuronal activity [19], to determine whether the reprogrammed neurons were active in the stroke-injured brain. At PSD 63, we observed YFP+NeuN+c-FOS+reprogrammed neurons (Figure 1g), indicating that reprogrammed cells express a marker of neural activity.

### Functional recovery is observed following reprogramming

We next asked if reprogramming astrocytes to neurons with *NeuroD1* was sufficient to promote functional recovery following sensory-motor cortical stroke. Motor behaviour was assessed using two measures of functional outcome: the grid-walking foot fault task, and Catwalk gait analysis.

The foot fault test was performed at PSD 4 (prior to reprogramming), PSD 28 and PSD 63 (21 and 56 days post-AAV injection). Stroke-injured mice displayed a significant motor deficit at PSD 4. By PSD 28, mice that received AAV5-*NeuroD1* were recovered in the foot fault task (Stroke_AAV5-*NeuroD1*, 0.22 +/- 0.53 slips; Stroke_AAV5-Cre, 1.58 +/- 0.26 slips; Sham_AAV5-Cre, 1.91+/- 0.37 slips; Uninjured controls, −0.15 +/- 0.34 slips; p<0.05; Figure 2a). As expected, uninjured and Sham_AAV5-Cre mice did not exhibit a deficit at PSD 4; however, after AAV-Cre injection, Sham_AAV5-Cre mice developed a transient motor impairment detected at PSD 28, suggesting that virus injection alone was sufficient to induce damage. Recovery was maintained in Stroke_*AAV5-NeuroD1* mice from PSD 28 until PSD 63; however, we observed spontaneous recovery in all groups with longer survival times, similar to previous reports [20,21]. To fully explore the long-term outcomes of the reprogramming strategy we used a more sensitive test where there was a detectable deficit in motor function long term.

**Fig. 2.**
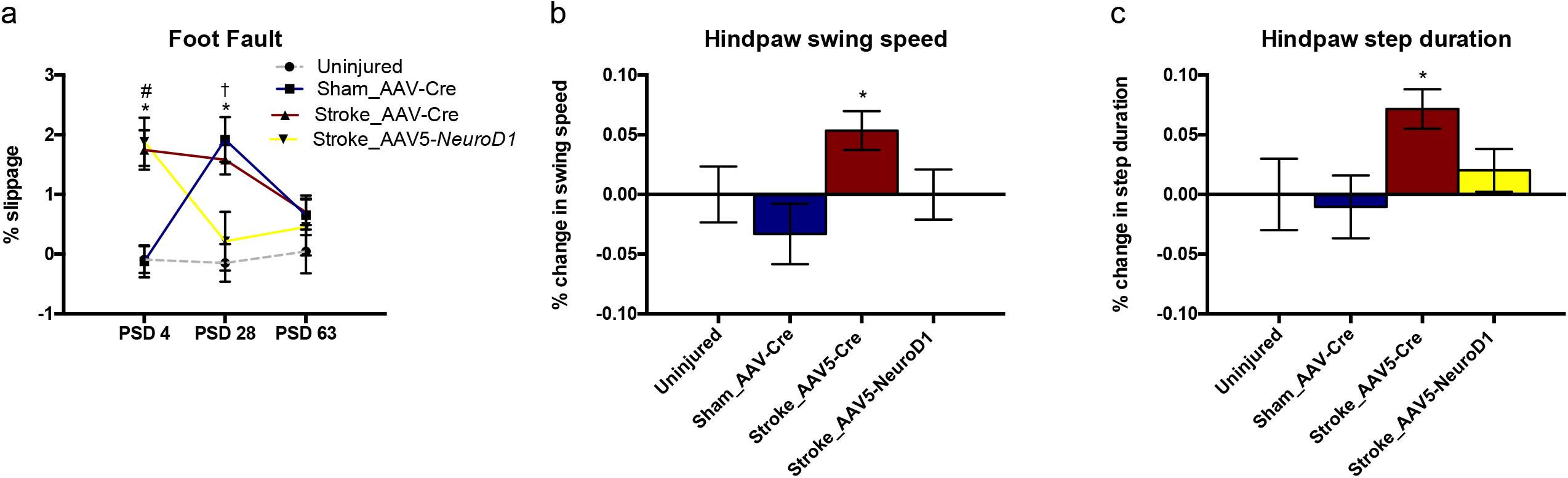
Functional outcomes associated with astrocyte-to-neuron reprogramming. a) Percent slippage in the foot fault test revealed a significant impairment in stroke injured mice prior to reprogramming (PSD 4) that was recovered by PSD28 in Stroke_AAV5-*NeuroD1* treated mice. A transient deficit was detected at PSD 28, following AAV-Cre injection into Sham controls. No long-term deficits were observed in any of the groups. Data are presented as mean ± SEM. *Stroke_AAV5-Cre (n=8) p<0.05 vs uninjured controls (n=6); #Stroke_AAV5-*NeuroD1* (n=7) p<0.05 vs uninjured controls; ^†^Sham_AAV5-Cre (n=11) p<0.05 vs uninjured controls. b) The duration of hindpaw swing speed and c) step duration were measured at PSD 63 by Catwalk digital gait analysis, normalized to uninjured control performance, and expressed as percent change between groups. Stroke-AAV5-Cre mice (n=26) displayed a significant impairment compared to Sham_AAV5-Cre controls (n=22) in both measures. Both measures improved in Stroke_AAV5-*NeuroD1* treated mice (n=30). *p<0.05 vs Sham_AAV5-Cre controls. Data are expressed as mean ± SEM.

Gait analysis at PSD 63 revealed deficits in Stroke_AAV5-Cre mice in measures of hindpaw swing speed and step duration (normalized to Uninjured control performance; p<0.05; Figure 2b-c). These deficits were attenuated in the Stroke_AAV5-*NeuroD1* mice (Figure 2b-c). Finally, we asked if functional recovery correlated with a change lesion volume. We observed no difference in stroke lesion volume in *AAV5-NeuroD1*- and AAV-Cre-injected mice (Stroke_AAV5-*NeuroD1*, 0.06 ± 0.03 mm^3^; Stroke_AAV-Cre, 0.07± 0.01 mm^3^; p=0.62) at PSD 63, when motor recovery was observed. Taken together, these findings suggest that astrocyte-to-neuron reprogramming is sufficient for improved motor function following stroke but with no concomitant change in lesion volumes.

## Discussion

Here we demonstrated that *NeuroD1*-mediated astrocyte reprogramming results in the production of mature, active and regionally appropriate neurons that survive long term and correlate with motor improvement following cortical stroke. Previous studies have demonstrated the feasibility of direct cellular conversion *in vivo* [7,9,12–14,22–25] but few have demonstrated improved behavioural outcomes following injury [12,17]. Using a foot fault task and gait analysis, we demonstrate that astrocyte-to-neuron reprogramming leads to functional improvement as early as three weeks post-reprogramming, and sustained long-term recovery. Our findings corroborate recent work in a mouse model of cortical ischemia [17] and lend support to the use of reprogramming with NeuroD1 as a regenerative strategy for stroke repair. Our studies were performed in a similar model of injury and showed similar timelines of improved motor function, as well as the production of both upper and lower layer cortical neurons. Interesting, while Chen et al. showed that a dual AAV9 system led to 80% reprogramming efficiencies (by 17d) and downstream motor gains, we showed only 20% (by 21d) is sufficient for long-term recovery. Together, our studies highlight the robustness and reproducibility of this Neuro-D1 based approach.

The endothelin-1 model of cortical injury produced reproducible lesions with small but significant motor deficits that were detectable using gait analysis until PSD 63 (the longest time examined), and lesion volumes similar to those reported previously [26,27]. Endohelin-1 is a commonly used preclinical model of ischemic stroke as it mimics the pathophysiological processes that are more closely related to the clinical condition, including excitotoxicity, generation of reactive oxygen species, related inflammatory responses, and gliosis [28,29]. Importantly, this model results in functional impairments can be recovered, which is critical for exploring the efficacy of cell therapy/regenerative interventions for stroke recovery. An important next step is to determine whether functional recovery can be achieved with increased tissue loss.

We found that functional improvement correlated with the generation of active (c-Fos expressing) cortical neurons that expressed appropriate layer-specific markers (CTIP2, TBR1 and CUX1). This supports recent findings using NeuroD1 [17] and a combination of the TFs *Nurr1* and *Ngn2* to reprogram astrocytes to neurons following a stab wound injury [7]. Moreover, this suggests that that local cues in the cortex may direct the generation of specific cortical subtypes in newly reprogrammed neurons following the initial acquisition of neuronal fate, similar to the production of regionally appropriate cell types from other cell sources as reported from both transplanted and endogenous stem cells [30–32]. Recent studies highlighting astrocyte heterogeneity raise the possibility that functional recovery could be due to the removal (via reprogramming) of subtypes of astrocytes that are detrimental to injury repair [33,34]. Specifically blocking reactive astrocytes (A1) was shown to be neuroprotective in a model of Parkinson’s Disease [35], and a recent study suggests that astrocyte reprogramming may specifically decrease this population [36]. Whether functional recovery is a direct result of the production of new neurons or due to the removal of astrocytes is an exciting question for future studies.

Interestingly, we show that reprogramming as few as 15% of the astrocytes in perilesional cortex (based on a transduction efficiency of 50% and reprogramming efficiency of about 30%) leads to behavioural improvement. Given that astrocytes play important roles in the normal and injured brain [37–41] and given the heterogeneity of this cell population, investigation into how the degree of astrocyte conversion impacts repair and recovery is crucial.

We chose to use AAV5 for our NeuroD1 delivery based on previous work suggesting enhanced astrocyte targeting [42]. A number of different viral delivery methods have been used for in vivo reprogramming (lentivirus, adenovirus, AAV9 [25]). These have shown varied transduction and reprogramming efficiencies and may preferentially transduce different astrocyte subtypes or other neural cell types, thus highlighting the importance of the gene delivery strategy.. In our study, we used an AAV5 strategy to drive NeuroD1 expression under the control of the GFAP promoter. We performed reprogramming at 7 days post-stroke in order to preferentially target cortical astrocytes and not neural stem cells that also express GFAP [43–46], as we have previously demonstrated that there is a rapid decline in the number of migratory subependymal zone (SEZ)-derived neural stem cells present in the cortex by this time point [16]. Further, we did not see transduced cells in the SEZ of reprogrammed mice. Previous work has shown a degree of non-specific expression from the GFAP promoter [7,47]. We used a short GFAP promoter with enchanced astrocyte specificity [47]. However, we still found a small percentage (~3-4%) of NeuN+YFP+ cells in our AAV5-Cre injected control group at both PSD 28 and 63 that may have resulted from a small degree of leakiness of the GFAP promoter or rare GFAP expression in neurons. Nonetheless, considering the significant functional improvement detected in AAV5-NeuroD1-treated animals, any spontaneous or injury-induced conversion provided negligible improvement. Consideration of reprogramming strategies will be an important next step for the application of this technology to the treatment of CNS injury and disease.

Our results show that reprogramming astrocytes into neurons leads to the generation of regionally appropriate neurons and is associated with functional recovery. Our work supports other recent studies investigating the outcome of reprogramming in the context of CNS injury and disease [7,12,17], and highlights the promise of reprogramming as a new regenerative strategy for stroke repair.

## Materials and Methods

### Mice

Adult (8-12 week old) male and female Rosa26-loxP-stop-loxP-YFP (YFP^fl^; Jackson Labs: B6.129X1-*Gt(ROSA)26Sortm1(EYFP)Cos*/J) Cre-conditional transgenic reporter mice were used in this study [48]. Mice were group housed in a barrier facility with a 12 hour light/12 hour dark cycle and allowed free access to food and water. Experiments were conducted according to protocols approved by the Animal Care Committee at the University of Toronto and performed in accordance with guidelines published by the Canadian Council for Animal Care.

### Endothelin-1 (ET-1) stroke

ET-1 (Calbiochem) was used to induce stroke as previously described [27,49]. Mice were anesthetized using isoflurane and mounted onto a stereotaxic apparatus. The scalp was incised and a small hole was drilled into the skull at the injection location. One μL of 400 pmol ET-1 dissolved in sterile PBS was injected into the sensorimotor cortex at +0.6 AP, +2.25 ML, −1.0 DV from bregma. A 26 gauge Hamilton syringe with a 45 degree beveled tip was used to inject ET-1 at a rate of 0.1 μl/min. The needle was left in place for 10 minutes after completion of the ET-1 injection to prevent backflow, then slowly withdrawn. Body temperature was maintained at 37°C using a heating pad, and animals were allowed to recover under a heat lamp. Ketoprofen (5.0 mg/kg; s.c.) was administered for post-surgery analgesia.

### Adeno Associated Virus (AAV) injections

AAV5 vectors containing NeuroD1-T2A-Cre or Cre driven by the GFAP promoter (gfaABC(1)D; [47]) were generated [NeuroD1AAV5-GFAP(0.7)^promoter(gfaABC(1)D)^-*NeuroD1*-T2A-Cre-WPRE (AAV5-*NeuroD1*; 5.0 x 10^12^ GC/mL) and AAV5-GFAP(0.7)^promoter(gfaABC(1)D)^-Cre-WPRE (AAV5-Cre; 1.1 x 10^13^ GC/mL)], and particles were packaged by Vector BioLabs (PA, USA). Seven days following stroke, mice were anesthetized with isoflurane and mounted onto a stereotaxic apparatus. One μl of AAV containing either *NeuroD1* or Cre was injected into the ipsilesional cortex at a rate of 0.1 μl /min at each the following coordinates: +0.6 AP, +2.25 ML, −1.0 DV; +1.6 AP, +2.25 ML −1.0 DV; and −0.4 AP, +2.25 ML, −1.0 DV mm from bregma. The needle was left in place for 10 minutes after each AAV injection to prevent backflow and then slowly withdrawn. Body temperature was maintained at 37°C using a heating pad, and animals were allowed to recover under a heat lamp. Ketoprofen (5.0 mg/kg; s.c.) was administered for post-surgery analgesia.

### Tissue processing, immunohistochemistry and quantification

Mice were anesthetized with an overdose of Avertin (250 mg/kg; i.p.), and perfused transcardially with saline followed by 4% paraformaldehyde (PFA). Brains were removed, post-fixed for 4 hours with 4% PFA then placed in 30% sucrose to cryopreserve. Brains were frozen and sectioned (20 μm) using a cryostat.

At the time of processing, sections were blocked with 10% normal goat serum (NGS) and 0.3% triton in PBS and labeled with primary antibodies (Table 1) in PBS overnight at 4°C, followed by incubation with secondary antibodies and a nuclear stain (Table 1) in PBS for 1 h at room temperature (RT). Colocalization analysis was performed by analyzing overlapping expression in 3-5 confocal stacks in 3 coronal sections per animal viewed at 20x magnification located at +1.6, +0.6, and −1.0 AP from bregma (Figure S1a). At each of these locations, imaging was performed at 3 regions of interest based on their distance from the infarct (defined as the heavily nucleated region devoid of NeuN+ staining) at 0-100 μm (within the lesion), 100-200 μm, and >200 μm, medially and laterally. Cortical layer-specific markers were imaged from upper and lower cortical layers (Figure S1b). Imaging was performed using Zen 2011 software and a Zeiss confocal microscope. Linear adjustments of contrast and brightness were made to micrographs using the respective microscope software or Photoshop using the levels function as follows: Figure 1c - *Cre* (linear brightness adjustment, green channel), *NeuroD1* (linear brightness adjustment, all channels); Figure 1d - *NeuroD1* (linear brightness adjustment, all channels); Figure 1f - CUX1 (linear brightness adjustment, all channels); TBR1 (linear brightness adjustment, red channel); CTIP2 (linear brightness adjustment, red channel); Figure 1g - c-FOS (linear brightness adjustment, all channels). Threshold intensity was set according to background signal detected in controls

**Table 1.**
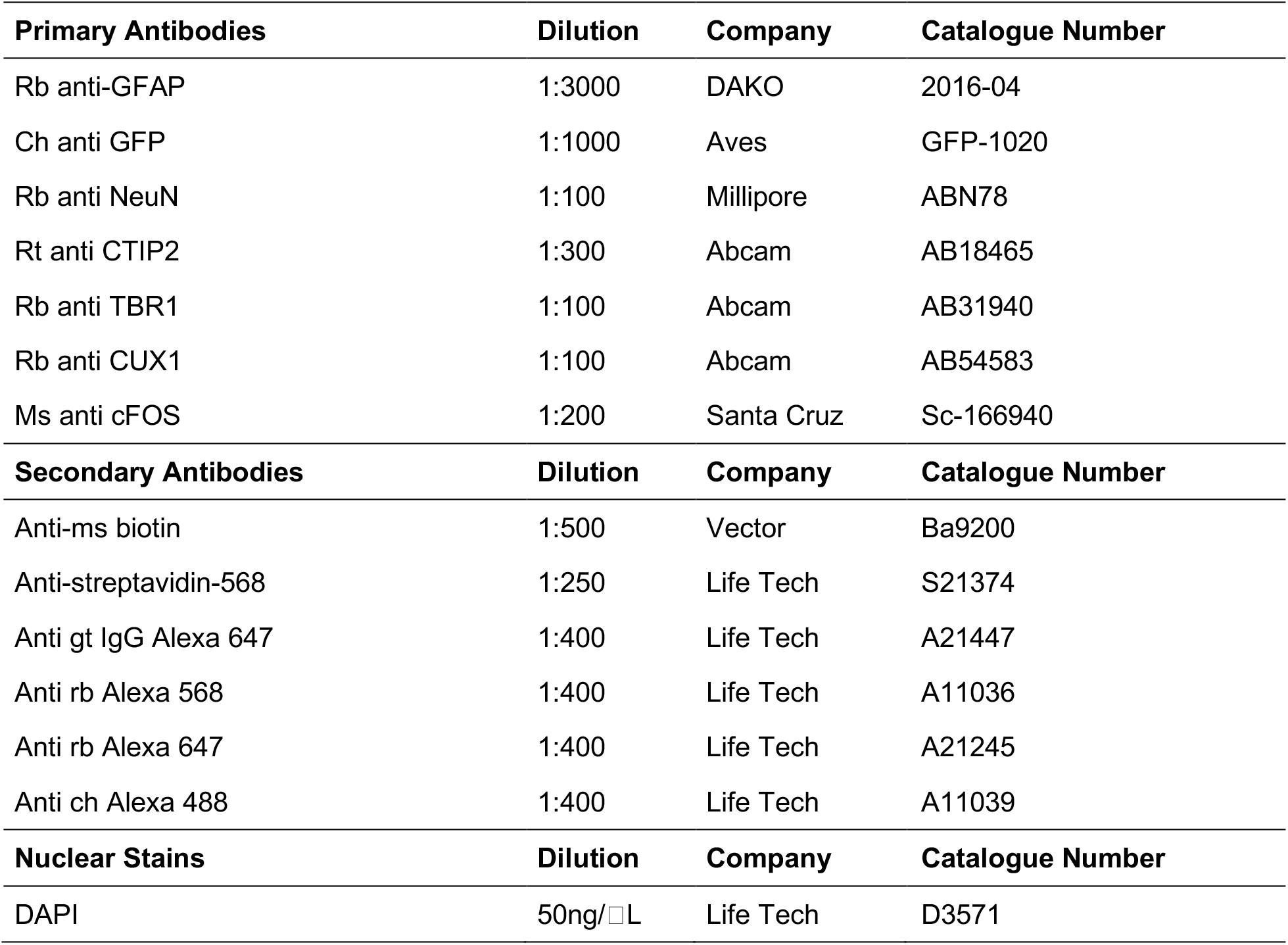
Summary of reagents used for immunohistochemistry

| **Primary Antibodies** | **Dilution** | **Company** | **Catalogue Number** |
| --- | --- | --- | --- |
| Rb anti-GFAP | 1:3000 | DAKO | 2016-04 |
| Ch anti GFP | 1:1000 | Aves | GFP-1020 |
| Rb anti NeuN | 1:100 | Millipore | ABN78 |
| Rt anti CTIP2 | 1:300 | Abcam | AB18465 |
| Rb anti TBR1 | 1:100 | Abcam | AB31940 |
| Rb anti CUX1 | 1:100 | Abcam | AB54583 |
| Ms anti cFOS | 1:200 | Santa Cruz | Sc-166940 |
| **Secondary Antibodies** | **Dilution** | **Company** | **Catalogue Number** |
| Anti-ms biotin | 1:500 | Vector | Ba9200 |
| Anti-streptavidin-568 | 1:250 | Life Tech | S21374 |
| Anti gt IgG Alexa 647 | 1:400 | Life Tech | A21447 |
| Anti rb Alexa 568 | 1:400 | Life Tech | A11036 |
| Anti rb Alexa 647 | 1:400 | Life Tech | A21245 |
| Anti ch Alexa 488 | 1:400 | Life Tech | A11039 |
| **Nuclear Stains** | **Dilution** | **Company** | **Catalogue Number** |
| DAPI | 50ng/□L | Life Tech | D3571 |

### Lesion Volume Analysis

Lesion volume was calculated from NeuN-stained sections, and defined as areas devoid of NeuN positive stain (Supplemental Figure 1a). Image J (National Institute of Health, USA) was used to measure this area in 5 x 20 μm thick coronal sections (200 μm apart) spanning the anterior-posterior extent of the injury site. This area was then multiplied by the distance between sections to estimate total infarct volume.

### Astrocyte Sorting and ImageStream Analysis

Mice were anaesthetized with isoflurane and cervically dislocated. The brain and meninges were removed and coronal slices were obtained using a scalpel blade. The cortex was collected from slices in the lesioned cortex, taking care to avoid the corpus callosum. The tissue was enzymatically and mechanically dissociated into a single-cell suspension using the magnetic activated cell sorting (MACS) Neural Tissue Dissociation Kit (Miltenyi Biotec 130-093-231). Myelin was removed using the MACS Myelin Removal Beads (Miltenyi Biotec 130-096-733). Dissociated cells were resuspended in 500 μL of PBS (azide and serum/protein free) and stained with a fixable cell viability dye (FVD eFluor 780; eBioscience, 65-0865-14) according to the manufacturer’s instructions. Cells were then washed with 10% FBS in PBS, resuspended in 500 μL of MACS buffer, stained with anti-GLAST-PE (Miltenyi Biotec 130-098-804), sorted using the MACS anti-PE MicroBeads UltraPure Kit (Miltenyi Biotec 130-105-639) according to the manufacturer’s instructions, and stained with Hoescht (1 μg/mL, BD Biosciences, 33342) for 30 min at RT, then resuspended in MACS sorting buffer.

Cells were analysed with an Amnis ImageStreamTM Mark II imaging flow cytometer and the resulting compensated image files were analysed using IDEAS analysis software (AMNIS). Single stained controls were collected for the compensation matrix. Focused, single cells were selected based on viability (exclusion of FVD e780 dye) and the presence of a nucleus (positive for Hoescht staining). Live nucleated cells were examined for PE-GLAST signal and YFP. The antibody binding patterns were confirmed in composite cell images of bright field and PE fluorescence generated from the ImageStream data.

### Foot Fault Analysis

Mice were placed onto a metal grid (1 cm spacing) that was suspended 12 inches above a table surface. Animals were allowed to walk around the grid for 3 minutes and were recorded from below. The number of steps and paw slips made with each forelimb were analyzed, and the difference in foot faults made was calculated as follows: [(#slips/#steps_ispilesional_) - (#slips/#steps_contralesional_)]. The following inclusion criteria were used: i) demonstrated deficit in stroke-injured animals or lack of deficit in uninjured animals (as +/- 2 standard deviations from the mean at post-stroke day 4 (PSD 4), ii) > 50 steps taken during the test.

### Gait Analysis

Prior to testing, mice were acclimated for at least 30 minutes in the behavioural suite. Gait analysis was performed using the automated Noldus CatWalk XT system (Noldus, Netherlands), according to manufacturer’s instructions. Briefly, mice were allowed to traverse a glass walkway illuminated with a light which measured a number of parameters regarding gait. A minimum of 3 successful trials was required, wherein the mouse took a direct path and speed did not vary more than 60% during the passage. Footprints were recorded using a high-speed video camera, and analyzed using CatWalk XT software across a range of variables surrounding paw placement and limb coordination while walking.

### Data Acquisition and Statistical Analysis

All experiments were conducted by an experimenter blinded to group allocation. Data was tested for normality using the D’Angostino and Pearson omnibus normality test or the Kolmogorov-Smirnow test. Unpaired two-tailed *t*-tests with Welch’s correction (for unequal variance) were used to compare two groups. One-factor analysis of variance (ANOVAs) followed by post hoc analysis (Tukey’s or Dunnet’s test) were used to compare 3 or more groups. Behavioural data were analysed using 2-way repeated measures ANOVA, followed by Bonferroni post-hoc tests when appropriate. Differences were considered significant at p < 0.05. Values are presented as mean ± SEM.

## Acknowledgments

The authors are grateful to Mira Puri for critical discussions and Ilan vonderwalde for assistance with behavioural analyses. ImageStream analysis was performed at the Lunenfeld-Tanenbaum Research Institute Flow Cytometry Core Facility.

## Author contributions

J.L. conducted experiments, analyzed, interpreted data and assembled data, and wrote the manuscript. T.L. conducted experiments and analyzed and interpreted data. C.P. conducted experiments, analyzed and interpreted data. E.D. analyzed, interpreted and assembled data. A.K. conducted experiments, analyzed and interpreted data. T.E. conducted experiments and analyzed and interpreted data. I.K. analyzed and interpreted data. K.W.A.B. conducted ImageStream experiments, analyzed and interpreted data. B.D. conducted experiments and analyzed and interpreted data. O.I. conducted experiments and analyzed data. N.S. conducted experiments and interpreted data. C.M. conceived and designed experiments, provided financial support, analyzed and interpreted data and wrote the manuscript. M.F. conceived, designed and conducted experiments, provided financial support, analyzed and interpreted data, assembled data and wrote the manuscript. All authors contributed to the final version of the manuscript.

## Additional information

### Data Availability

The datasets generated during and/or analysed during the current study are available from the corresponding author on reasonable request.

---

**Fig. S1.** Quantification of reprogrammed cells. a) Images were analyzed from 100 x 100 μm counting boxes in 3 sections per animal (+1.6, +0.6, −1.0 mm AP from bregma) in the area of the lesion (dashed line; defined as the heavily nucleated area devoid of NeuN staining), and in two adjacent regions of interest (up to a total of 200 μm) medially and laterally from the lesion border. b) Cortical layer-specific marker expression was analyzed within the areas of the red boxes (100 x 100 μm) from the same 3 sections.

