## Supplementary figures and images for "Direct reprogramming of astrocytes to neurons leads to functional recovery after stroke"

### Supplemental Figure S1

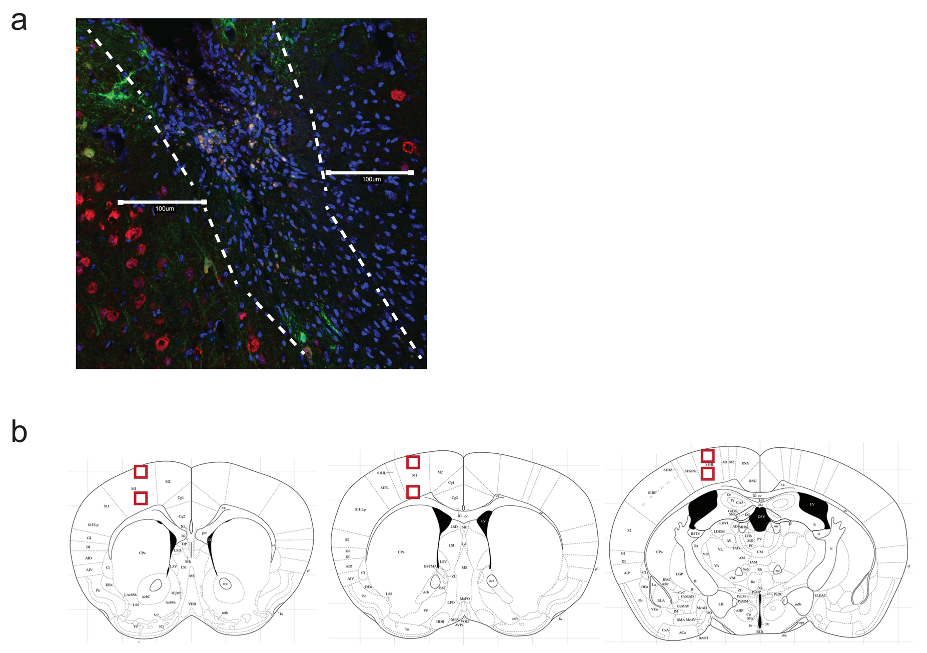
